# Decoding with multivariate pattern analysis is superior for optically pumped magnetometer-based magnetoencephalography compared to superconducting quantum interference device-based systems

**DOI:** 10.1101/2025.10.14.681912

**Authors:** Jiawei Liang, Yulia Bezsudnova, Anna Kowalczyk, Yu Sun, Ole Jensen

## Abstract

**Background:** Multivariate pattern analysis (MVPA) has become an increasingly important method for decoding distributed brain activity from neural electrophysiological recordings by leveraging both temporal and spatial features. These multivariate approaches have proven important for both cognitive neuroscience and brain-computer interfaces. MVPA might benefit from magnetoencephalography (MEG) systems based on optically pumped magnetometers (OPMs), as these sensors can be placed closer to the scalp, providing higher spatial resolution compared to conventional MEG systems that rely on superconducting quantum interference devices (SQUIDs). As OPM-based MEG systems become available at more institutions, it is essential to experimentally compare their performance with traditional SQUID-based systems using MVPA.

**Methods:** We adapted a visual object-word paradigm from a previous study, originally implemented on a TRIUX MEGIN SQUID system, to the FieldLine HEDscan OPM system. Participants were recruited and we recorded their ingoing brain activity while did the same task in both systems. Visual stimuli of different objects were presented alternately in two modalities: pictures and the corresponding written words. For each modality, MVPA was used to classify the objects from OPM and SQUID magnetometers data respectively. To further investigate the advantages of OPM, we evaluated the effect classification accuracy of two spatial factors by controlling the number of sensors included and the spatial frequency content of the sensor data.

**Results:** We found higher time-resolved decoding accuracy for the OPM compared to the SQUID data. Moreover, OPMs show higher classification performance compared to SQUIDs when controlling for the same number of sensors; consistently, the OPM system required fewer sensors to reach the performance limit of the SQUID system. Our analysis considering the spatial frequency content of the signal revealed that decoding accuracy plateaued for the SQUID system at lower spatial frequencies while the performance of the OPM system continued to improve when higher-order spatial components were included.

**Conclusion:** Our OPM-MEG system outperformed the SQUID-MEG system on MVPA on decoding of visual processing. This advantage of OPM is driven by its higher spatial resolution, resulting from the sensors being positioned closer to the head and thus able to capture higher spatial frequency components of the brain signal. OPM may facilitate cognitive neuroscience research as well as brain-computer interfaces by providing higher sensitivity when employing paradigms using multi-variate data analysis.

## Introduction

Multivariate pattern analysis (MVPA) has proven to be an invaluable tool across a wide range of cognitive neuroscience domains (Haxby et al., 2001; Haynes & Rees, 2006; Pereira et al., 2009; Cichy et al., 2014; Grootswagers et al., 2017; Guggenmos et al., 2018; Peelen & Downing, 2023; Wu et al., 2024) due to its ability to capitalize on both the spatial and temporal structure of brain activity (Bezsudnova et al., 2024; Cichy et al., 2016; Dirani & Pylkkänen, 2023; Stokes et al., 2015). The multivariate approach is also applied in brain-computer interfaces (BCIs) (Mellinger et al., 2007; Valente et al., 2019; Waldert et al., 2008; Wittevrongel et al., 2021) including novel brain-to-text efforts (Boyko et al., 2024; Défossez et al., 2023; Lévy et al., 2025). Compared to a cryogenic magnetoencephalography (MEG) system based on superconducting quantum interference devices (SQUIDs), optically pumped magnetometers (OPMs) allow for a more flexible sensor arrangement and smaller distances from the sensors to the scalp, which result in higher recorded brain signals (Brookes et al., 2022), theoretically improved spatial resolution (Iivanainen et al., 2017; Nugent et al., 2022), and the potential for more anatomically tailored sensor coverage (Beltrachini et al., 2021), particularly in regions like the frontal lobes that are traditionally challenging to access with SQUID systems (Safar et al., 2024). These advantages over SQUID make OPM especially well-suited for studying spatially distributed brain processes via MVPA (Labyt et al., 2022; Wens, 2023). The potentially finer spatial resolution of OPM system may also help improve the limited performance of MEG reported in a related study (Banville et al., 2025; Benchetrit et al., 2024). Furthermore, since the OPM sensor array can be adapted to different head size across the life-span, it sets the stage for developmental studies using MVPA in children, infants and possibly even foetuses (Corvilain, Capparini, et al., 2025; Corvilain, Wens, et al., 2025; Feys et al., 2022).

### QUESTIONS OF HYPOTHESIS

Although OPM sensors detect stronger neural signals, they exhibit higher levels of intrinsic noise compared to SQUID-based systems (Iivanainen et al., 2019). However, this limitation may be less critical in the context of MVPA, because classifiers rely on pattern differentiation across multivariate spaces and may remain robust in the presence of noise (Grootswagers et al., 2017). Therefore, we hypothesize that OPM systems can attain a performance gain over SQUID from the greater magnitude of the neural signals captured by the sensors closer to the scalp. Moreover, this advantage should be more pronounced in signal components of higher spatial frequency, as the amplitude decrease exponentially with the order of spherical expansion signals in the MEG sensor array (Taulu, 2008, p. 41).

Furthermore, it remains to be examined how the number of MEG sensors affects the MVPA performance in our specific experiment. Spatial-frequency analyses of MEG/EEG indicate that sensor count directly governs the range of spatial frequencies that can be sampled (Iivanainen et al., 2021).

### HOW TO ANSWER

To evaluate the performance of OPM-MEG in multivariate analysis, we here empirically test the efficacy of MVPA using SQUID and OPM data. Our paradigm includes two modalities: pictures and the corresponding written words (Fig 1). The data were analysed using pairwise decoding (Cichy et al., 2014). By directly assessing decoding performance across different modalities and conditions with controlled experimental paradigm and preprocessing pipeline, we aimed to determine whether and how the robustness and spatial advantages of OPMs translate into measurable improvements in neural decoding.

**Figure 1.**
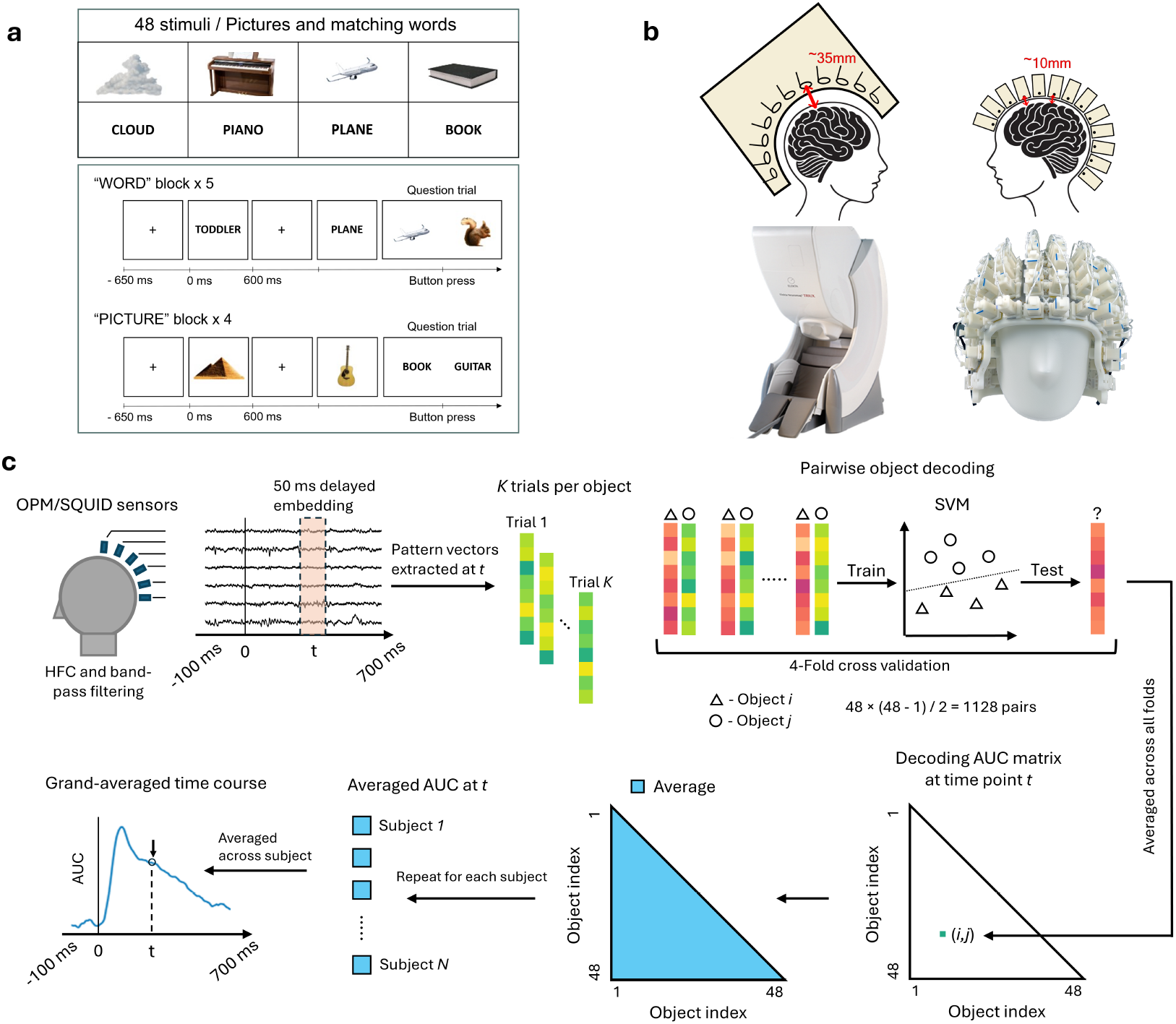
Experimental design and multivariate analysis. (a) Experiment paradigm. The stimulus set consists of 48 pictures and corresponding words in text. Five blocks for word stimulus and four for picture were arranged alternately. Within each block, each stimulus was presented in random order and repeated for 4 times, with one time followed by a question trial in the opposite modality. The stimulus duration was 600 ms and the fixation duration was randomly 550 – 750 ms. (b) Demonstration of the MEGIN Elekta SQUID system (left) and FieldLine HEDscan OPM system (right). (c) MVPA pipeline of pairwise object decoding. Epochs were delimited 100 ms before and 700 ms after the stimulus onset from pre-processed OPM/SQUID recordings, yielding 16 trials for each picture and 20 for each word. For each timepoint, sensor space pattern vectors were extracted with 50 ms time-delayed embedding. For each of the total 1128 object pairs, a binary SVM classifier was trained and evaluated via 4-fold cross validation. The decoding accuracy was averaged across all conditions and participants at each time point, which ultimately produced the AUC time course for picture and word modalities.

## Methods

### Experimental paradigm

The experiment paradigm was adapted from a previous study (Iamshchinina et al., 2022), with some stimuli replaced and the task slightly modified (Fig 1a). Visual stimuli of 48 objects were presented in the form of pictures and the corresponding written words (font: Arial, bold, 75). The stimulus measured 400 × 400 pixels and subtended a visual angle of 6°, which was projected on a screen with grey background. Projectors of identical model (VPixx PROPixx) were used in OPM and SQUID sessions both operating at 120 Hz refresh rate.

There were nine blocks in the experiment (five blocks for words and four blocks for pictures). Within each block, the participant viewed a series of stimuli for 600 ms each, which were separated by fixation crosses for a random duration of 550 to 750 ms. There were randomly inserted questions on average every fifth trial that asked the participant to choose the last seen stimulus from two options presented in the opposite modality of that block. These questions were included to keep participants attentive, and responses were made via button press within a timeout of 3.5 s. Participants were allowed to take self-paced breaks between blocks, ranging from at least 5 s to a maximum of 300 s.

### Experiment participants

Twelve healthy adults participated in the study. Three participants were excluded due to data quality issues: one associated with OPM trigger failure, one with an interrupted OPM recording, and one showing abnormal noise patterns in the SQUID PSD. Finally, nine participants (mean age = 25.1 +/-3.1 years old; 6 female) were included in consequent analysis. The order in which participants underwent OPM and SQUID sessions was counter balanced. All participants were native English speakers with normal or corrected-to-normal vision.

The study was conducted at the Centre for Human Brain Health in Birmingham at the University of Birmingham. The study was approved by University of Birmingham Ethics Committee. The participants provided written informed consent and received a £15 per hour compensation for their participation.

### MEG Acquisition

SQUID data were acquired using a 306-sensor TRIUX MEGIN system (Fig 2b, left), consisting of 204 orthogonal gradiometers and 102 magnetometers. An online band-pass filter from 0.1 to 330 Hz was applied before the data were sampled at 1000 Hz.

**Figure 2.**
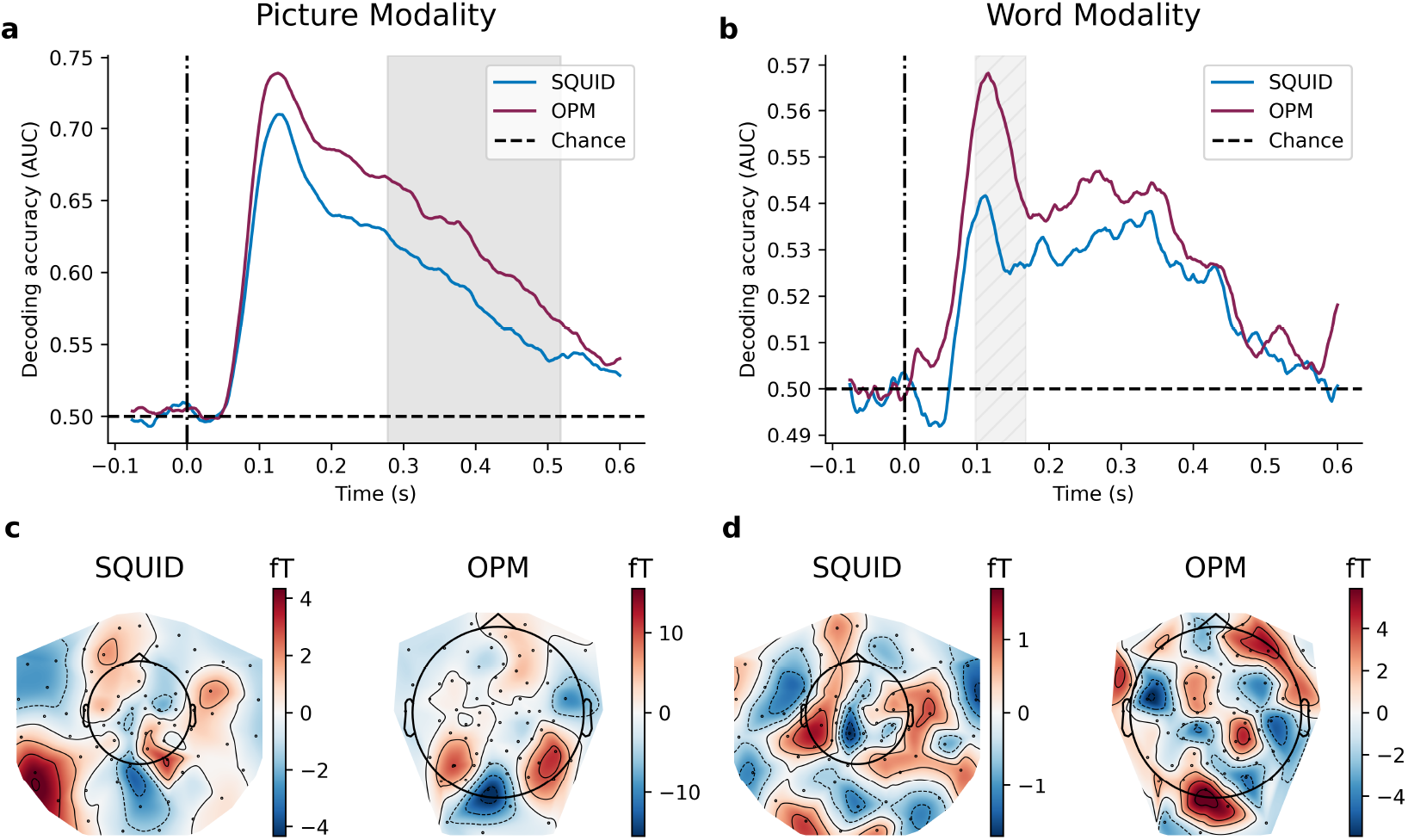
Time-resolved multivariate pairwise object decoding. (a)(b) Time-resolved decoding AUC for picture and word modalities with 68 MEG sensors. Purple line, OPM; blue line, SQUID; dashed line, chance level. The highlighted area denotes the significant cluster (*p* <.05) where AUC of OPM was higher for picture modality. The hatched area represents a cluster (*p* =.057) where AUC of OPM was higher for word modality. Cluster-based permutation test was one-tailed and was controlled for multiple comparisons over time. (c)(d) Topographical activation patterns at peak decoding times (127 ms for pictures; 114 ms for words) derived from classifier weights (Haufe et al., 2014), averaged across participants, stimulus objects and time points of delayed embedding.

OPM data were recorded with a FieldLine HEDscan system (Fig 2b, right) with 73 single-axis SERF-OPM sensors operating at 5000 Hz sampling rate with an online low-pass filter at 500 Hz. The OPM sensors were uniformly distributed in the helmet, covering the whole head. One sensor was used as eyeblink detector. A few sensors randomly failed or suffered from high noise level, which were disabled before recording.

### Data Pre-Preprocessing

The data were analysed using the open-source toolbox MNE Python v1.7.1 (Gramfort et al., 2013) following the standards defined in the FLUX Pipeline (Ferrante et al., 2022). A two-step sensor dropping method was applied to OPM data at start. First, the sensors with excessive broad-band noise in the raw power spectral density (PSD) were visually identified and rejected. Subsequently, a homogeneous field correction (HFC; order=2) filter (Tierney et al., 2021) was tentatively applied. Any sensor that maintained an abnormal PSD afterwards was further excluded. This resulted in an average of 69.8 sensors (a minimum of 68 sensors). Finally, unprocessed raw data were retrieved from the remaining sensors for the following steps.

To normalize the number of sensors across both systems and participants, SQUID magnetometers and OPMs were subsampled to a maximum of 68 (and fewer, as described in the spatial property analysis section). Subsampling was performed using random k-means clustering in the sensor-layout space, maintaining even coverage of the helmet throughout the process. After that, an HFC filter (order=2) was applied to the subset of MEG sensors. The data was further low pass filtered at 200 Hz and then down sampled to a sampling rate of 500 Hz. A notch filters at 50 Hz and 100 Hz were used to remove line noise. A band-pass filter between 1 Hz to 100 Hz was applied to reduce low frequency fluctuations introduced by lift operation nearby and other high frequency noise. The paradigm events were annotated based on the trigger channel information.

### Pairwise object decoding

As shown in Fig 1c, the MEG data acquired with SQUID and OPM systems were first segmented into trials (-100 to 700 ms) that were time-locked to the stimulus onset. All the N sensor values were further concatenated along a sliding time window of ±25 ms (25 time points in total), e.g. resulting in a N × 25-dimensional feature vector for each time point reflecting the N sensors.

MVPA was applied to the processed MEG data to decode the object identity pairwise, separately for each participant’s OPM and SQUID data. First, the feature vectors associated with all trials of each object in the paradigm were extracted, yielding 16 trials per picture and 20 trials per word. For each stimulus modality, binary classification of object pairs was performed at each time point using a support vector machine (SVM) with four-fold cross-validation, chosen so that the number of trials was evenly divisible by the folds. Within each fold, features were z-scored to zero mean and unit variance prior to classification. Decoding performance was quantified using the area under the curve (AUC) averaged across all folds.

For each participant and MEG system, the aforementioned steps resulted in the strictly lower triangular part of a 48 by 48 representational dissimilarity matrix of the pairwise object decoding at each time point. The decoding AUC was further averaged across all object pairs (n = 1128) and participants (n = 9) at each time point, ultimately resulting in AUC time courses for picture and word modalities.

To investigate the spatial distribution of brain activity underlining the decoding, the topographical activation patterns were extracted using back-projection of classifier weights via the method described in (Haufe et al., 2014).

### Spatial sampling and spatial frequency analysis

To evaluate the SQUID and OPM systems, we first subsampled the sensors from 10 to 68 and repeated the preprocessing and MVPA decoding pipeline for each sensor count. The averaged decoding AUC over time was then calculated as a function of the number of sensors.

Complementary to this sensor-count analysis, we also examined the contribution of signal components at different spatial frequencies. To investigate this, we replaced the HFC algorithm with adaptive multipole models (AMM) (Tierney et al., 2024) using spherical harmonics. Since higher harmonic orders represent higher spatial frequency components of the modelled brain signal (Nortje et al., 2015; Tierney et al., 2022), we progressively increased the number of inner harmonics included in the reconstruction and then repeated the analysis pipeline. A maximum expansion order of 7 was imposed, constrained by the 68 sensors available (Tierney et al., 2022) and the external harmonic order was fixed at 2 to match the order used for HFC in the previous analysis.

A separate analysis was further performed by fitting individual functions from the inner orders of the spherical expansion to the decoding AUC, yielding the curves shown in Fig 4. A three-parameter sigmoid function was adopted as follows:

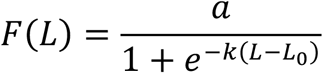

where ***L*** denotes the maximum order of harmonics, ***k*** the steepness of the function, ***a*** the upper limit and ***L***_***0***_ the middle point. A larger ***k*** indicates that decoding performance increases rapidly with lower spatial frequency components and saturates with higher frequencies, whereas a smaller ***k*** indicates greater reliance on higher-frequency components.

### Statistical Analysis

To control for multiple comparisons over time-points when compared the classification results, a non-parametric cluster-based permutation test (Maris & Oostenveld, 2007; Sassenhagen & Draschkow, 2019; Smith & Nichols, 2009) was applied. A cluster-forming threshold corresponding to *p* <.05 (one-tailed) was applied at the sample level. Cluster-level significance was assessed against a one-tailed null distribution of the maximum cluster statistics obtained from 10000 random sign-flip permutations, and clusters with *p* <.05 were considered significant. To evaluate the differences in the growth trends of decoding AUC with harmonic order between OPM and SQUID data, a Wilcoxon signed-rank test was conducted to compare the fitted steepness ***k*** between the two systems. Fits with an adjusted *R*^2^ below 0.8 were excluded from statistical testing as outliers.

## Results

We conducted a within-participant comparison of MEG signals acquired with a conventional SQUID system and a single-axis OPM system, with MVPA applied to a visual object and word paradigm (Fig 1a). Participants responded accurately to the attention-check questions (OPM: M = 94.5%, SD = 5.40%; SQUID: M = 96.5%, SD = 2.35%). A Wilcoxon signed-rank test revealed no significant difference in accuracy across MEG systems (*V* = 15, *p* =.43), indicating that participants maintained comparable levels of attention throughout the task.

The MEG data were preprocessed and time-resolved object decoding accuracy (AUC) for each modality was quantified. To further assess spatial properties, we systematically varied the number of sensors included in the analysis and examined the contribution of different spatial frequency components of the measured signals using a spherical expansion approach. The following sections present the results of these analyses.

### Time-resolved AUC of pairwise object decoding

The MVPA approach yielded a decoding AUC time course of each pair of visual stimuli for each participant. A grand average curve was computed across all object pairs. Then these grand averages were aggregated across participants to assess the comparative performance of the OPM and SQUID based MEG systems for the picture and word modalities.

We observed robust decoding for both pictures and words and with both OPM and SQUID systems. The overall profiles and value ranges of the AUC time course are in agreement with the findings of previous MVPA studies involving pictures and written or spoken words as stimuli (Iamshchinina et al., 2022; Dirani & Pylkkänen, 2023, Dirani & Pylkkänen, 2024). As exemplified by the full 68-sensor case (Fig 2a and b), the temporal patterns appeared similar between OPM and SQUID, whereas OPM exhibited higher AUC values across the predefined interval of effective decoding. This difference was most pronounced in the 278-518 ms window for pictures (*p* =.020, cluster permutation), and a trend was observed in the 98-168 ms interval for words (*p* =.057, cluster permutation).

The activation topographic maps were computed at the midpoint between the peak AUC latencies of SQUID and OPM for each modality to visualize the spatial patterns underlying decoding. The sensors contributing most strongly to picture-object decoding were predominantly located over occipital areas for both OPM and SQUID (Fig 2c), whereas for word stimuli the contributing sensors showed a more dispersed distribution across parietal, temporal, and frontal areas (Fig 2d). To account for participant-specific adjustment of OPM sensor positions, the average topographic maps were derived through clustering and interpolation across all participants’ sensor layouts.

Together, these results show that both systems produced robust decoding consistent with prior MVPA studies, with OPM showing systematically higher accuracy. We next examined how this advantage depends on sensor number and spatial frequency content.

### Number of sensors

We then repeated the MVPA considering a subsampling of the number of sensors from 10 to 68. The averaged decoding performance was computed within predefined time windows, 75–600 ms for pictures and 75–440 ms for words relative to stimulus onset, based on previous reports (Cichy et al., 2014; Dirani & Pylkkänen, 2023, Dirani & Pylkkänen, 2024). Fig 3a and b illustrate the averaged AUC value as a function of the number of sub-sampled sensors for picture and word modalities respectively. The MVPA performance increased from 10 to ∼30 sensors and plateaued after ∼40 sensors. We observed consistently higher AUC for OPM compared SQUID MEG which was most pronounced from 34-52 sensors for pictures (*p* =.033) as well as 24-30 and 34-68 sensors (respectively *p* =.035; *p* =.008) for words. In summary, decoding performance saturated with around 40 sensors, and OPM consistently achieved higher accuracy than SQUID, reaching comparable performance levels with fewer sensors.

**Figure 3.**
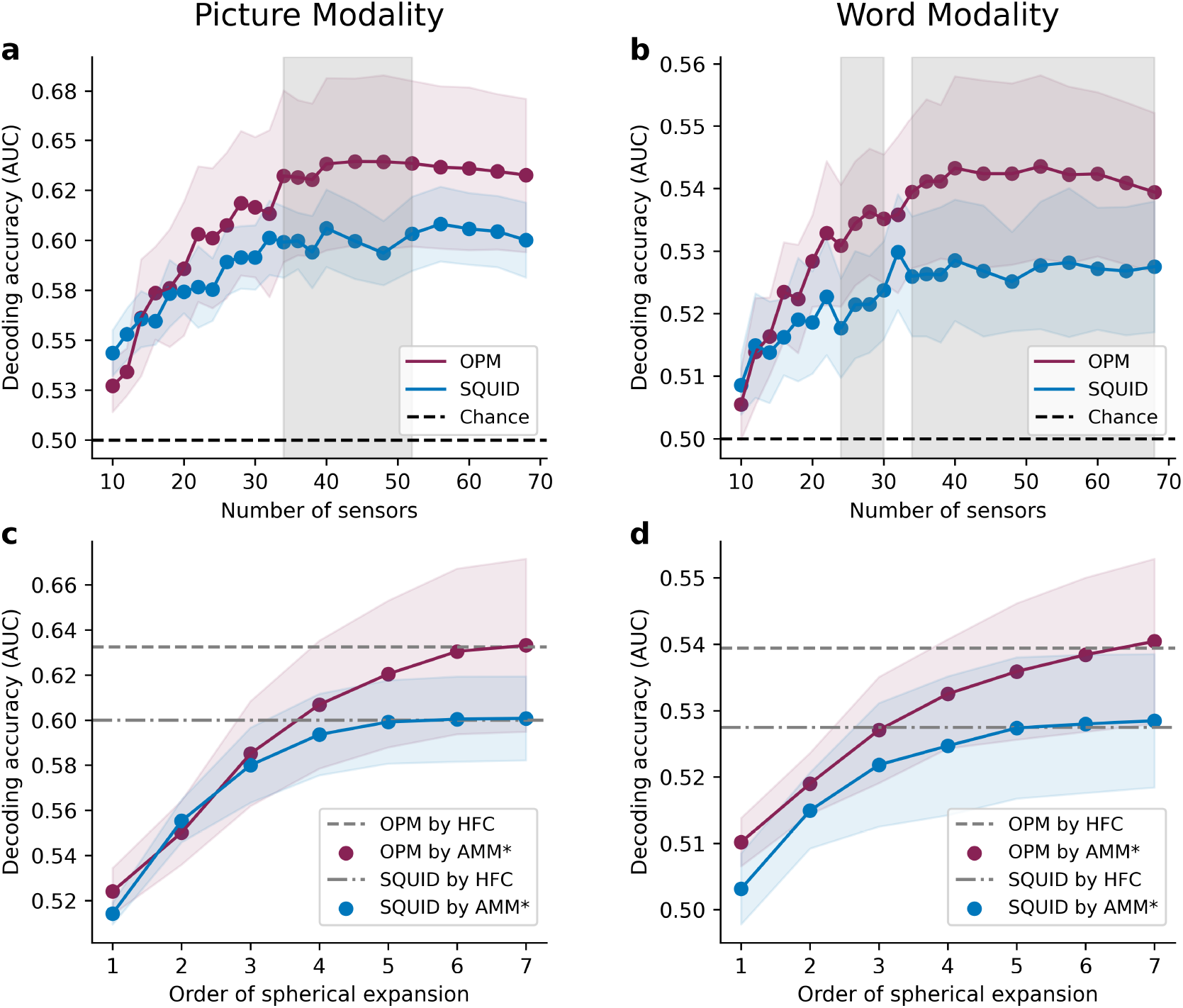
MVPA decoding performance as a function of the number of sensors and the order of spherical expansion. (a)(b) Mean decoding AUC (across 75–600 ms for pictures; 75–440 ms for words) with different numbers of sensors. Purple line, OPM; blue line, SQUID; dashed line, chance level (0.50). Shaded areas represent the 95% confidence interval (CI) across participants. The highlighted areas denote significant clusters (*p* <.05, one-tailed, controlled for multiple comparisons over sensor count) where AUC of OPM was higher than that of SQUID. (c)(d) Mean decoding AUC with the maximum order of spherical harmonics included. Coloured lines represent results obtained using the customized AMM filtering (shaded areas for 95% CI). Dashed and dash-dotted lines indicate the AUC obtained via the HFC algorithm, serving as reference levels for OPM and SQUID respectively.

### Spatial frequency components

Next, we examined the accumulative contribution to MVPA performance considering the spatial frequency components in the MEG signals. Fig 3c and d show the decoding AUC as a function of the order of spherical expansion using AMM filtering. The AUC from data pre-processed using HFC is presented as a reference where the inner spatial components were not limited. For the picture modality, OPM and SQUID MEG data exhibited comparable increase in decoding performance using the spherical harmonics from 1 to 3. While the AUC of SQUID plateaued at order 5, the OPM performance improved until order 6. For the word modality, the performance was better for OPMs compared to SQUID and this advantage tended to increase as higher order harmonics where included.

We next examined the spherical expansions for each participant. As shown in Fig 4a-d, the data were generally well fitted by the three-parameter sigmoid function (adjusted *R*^2^ > 0.9 for most participants), except for one participant whose parameter estimate was constrained at the bound for word modality, yielding low goodness-of-fit (*R*^2^ = 0.31 for OPM and 0.74 for SQUID). This participant was therefore excluded from statistical testing for words. Fig 4e and f show the group difference in steepness ***k*** between OPM and SQUID. OPM exhibited significantly lower steepness than SQUID for both pictures (*V* = 0, *p* =.004) and words (*V* = 2, *p* =.023), reflecting a more gradual increase that extended into higher-frequency components.

**Figure 4.**
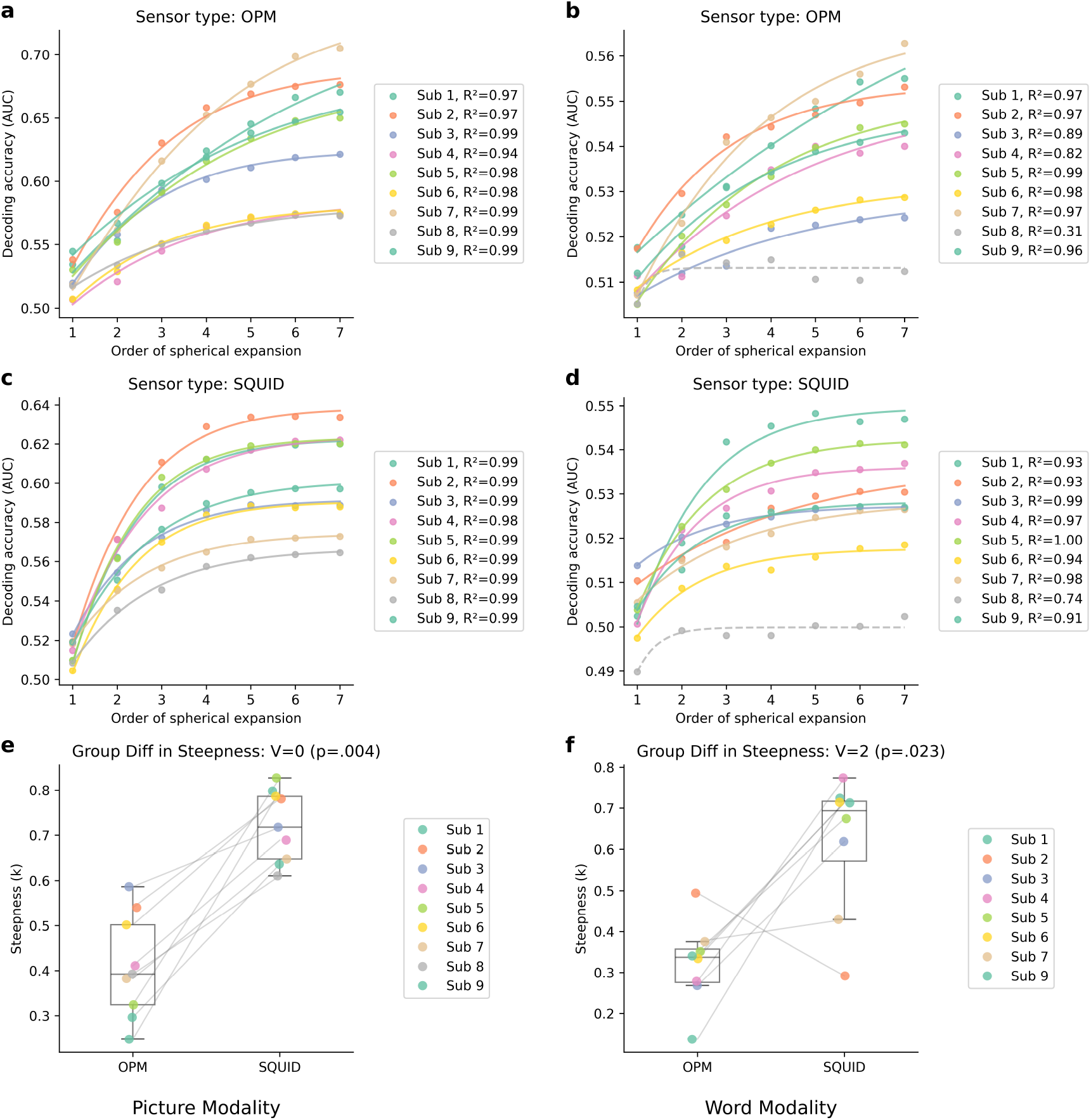
Individual decoding performance and sigmoid fits across orders of spherical expansion. Decoding accuracy (AUC) as a function of spherical harmonic expansion order of OPM (a)(b) and SQUID (c)(d) signals for the picture and word modalities, respectively. Data from individual participants (coloured dots) were fitted with a three-parameter sigmoid function (solid lines). Adjusted *R*^2^ values indicated good model fits across participants, except for one outlier in the word modality (OPM: *R*^2^ = 0.31; SQUID: *R*^2^ = 0.74), which was excluded from group-level analysis for that modality. (e)(f) Boxplots show the distribution of fitted steepness parameter ***k*** across participants, with paired individual data points overlaid. Steepness was significantly lower for OPM compared to SQUID in both picture (Wilcoxon signed-rank test, *V* = 0, *p* =.004) and word modalities (*V* = 2, *p* =.023).

Taken together, the findings indicate that OPM decoding continued to benefit from higher-order spatial components, whereas SQUID performance saturated earlier, suggesting that OPM captures finer spatial details of the neural signal.

## Discussions

In this study, we compared the performance of multivariate-pattern analyses from an OPM and a SQUID MEG system in terms of discriminating different visual stimuli of pictures and different words. We found that our OPM outperformed the SQUID MEG system in general (Fig 2a and b). The advantage of OPM remained evident when we sub-sampled the sensors and it attained the plateau performance of SQUID with much fewer sensors (Fig 3a and b). Furthermore, we studied how the decoding accuracy scaled with the spatial frequency content of the signal being decoded. The decoding accuracy plateaued for SQUID at lower spatial frequencies while OPM achieved a performance gain from higher-order spatial components (Fig 4).

The AUC time course revealed a sustained advantage of OPM over SQUID throughout the decoding interval (Fig 2a and b). The decoding peak occurred at similar latencies for both systems; however, OPM maintained a clear advantage even at the peak time. This effect was particularly pronounced in the word modality, where the difference in AUC between OPM and SQUID was greatest at the peak. These findings highlight two aspects of OPM’s superiority: a sustained decoding advantage across the entire decodable interval, which enhances sensitivity to the temporal dynamics of sensory processing; and an increased peak decoding capacity, which is particularly relevant for brain–computer interface applications that exploit the strongest representational windows.

Whereas previous simulation studies based on spatial sampling and source localization analyses (Iivanainen et al., 2021; Tierney et al., 2020) suggested that ∼90 sensors or more may be needed to adequately sample neuromagnetic signals under typical MEG paradigms, our subsampling analysis indicated that the performance of object decoding plateaued around 40 sensors for both OPM and SQUID (Fig 3a and b). Importantly, the gap between SQUID and OPM could not be compensated by incorporating more sensors. Moreover, when the number of sensors must be reduced, the decline in MVPA performance will not appear as fast. Remarkably, even with fewer than 30 sensors, OPM still achieved decoding performance comparable to the upper bound of a SQUID system, demonstrating its potential to facilitate wearable and affordable BCI systems where the number of sensors is constrained (Fedosov et al., 2025; Marhl et al., 2025). Further work is needed to determine whether this advantage of OPM generalizes to source localization or other MEG analysis approaches.

Our analysis of the relationship between order of spherical expansion and decoding performance further confirmed the source of the advantage of OPM (Fig 3cd, Fig 4). At equivalent sensor densities, OPM benefited more from higher-order spatial components than SQUID did. This provides experimental support for the theoretical prediction that the amplitude coefficients of spherical harmonics decay with distance ***r*** according to *r*^−(*l*+2)^ (Taulu, 2008, p. 41), where ***l*** is the harmonic order. Thus, sensors placed closer to the scalp capture stronger signals, and the relative gain becomes even larger at higher orders. In summary, the closer sensor proximity in OPM yields an amplitude boost that is greater for high spatial frequency components, enabling OPM to leverage them effectively despite its higher noise level.

When comparing the picture and word modalities, we first observed that decoding AUC was substantially lower for the word modality. This difference was also reflected in the topographic maps of decoding activation averaged over all participants, where word-related responses showed lower amplitudes (Fig 2c and d). By contrast, the relative advantage of OPM over SQUID was more pronounced in the word modality. This was manifested as a sharper decoding peak (Fig 2b), as well as clearer AUC differences emerging even with fewer sensors (Fig 3b) or at lower orders of spherical expansion (Fig 3d). These results suggest that OPM retains sufficient sensitivity to capture meaningful representations even when the overall neural signal is weaker. Consequently, OPM may offer particular benefits in research topics where signal strength is expected to be low, such semantic processing, memory reactivation, or other higher-order cognitive functions.

Several limitations of the present study should be noted. First, our OPM system employed single-axis sensors rather than dual-axial or tri-axial ones. Multi-axial OPM sensors are becoming increasingly available (Boto et al., 2022; Rea et al., 2022), and they may further facilitate noise reduction for OPM (Brookes et al., 2021; Tierney et al., 2022) and possibly also contribute to better multivariate decoding. Future work should directly evaluate whether the advantage of OPM over SQUID generalizes or even amplifies with multi-axial configurations. Second, our analysis focused exclusively on object-level decoding and did not examine higher-order semantic representations due to small number of participants. While OPM demonstrated a clear advantage in distinguishing visual objects, it remains to be determined whether this advantage extends to more abstract decoding tasks, such as semantic category or conceptual processing. Third, the current paradigm was restricted to the visual modality. Extending OPM-based MVPA to other domains, such as auditory processing, language, or motor actions, would provide a more comprehensive assessment of its advantages over SQUID. The study included nine participants. While this provides adequate power to observe large effects in the MVPA comparison between OPM- and SQUID-MEG, increasing the sample size would improve sensitivity to nuanced patterns, refine effect-size estimates, and enhance generalisability.

In conclusion, our findings provide strong experimental evidence that OPM-MEG offers superior performance over SQUID-MEG in multivariate pattern analysis of visual processing. We experimentally show that our OPM system has higher spatial resolution compared to our SQUID system. While much of the recent discussion on OPMs has emphasized their potential for improved source modelling (Tierney et al., 2020; Beltrachini et al., 2021; Nugent et al., 2022; Sun et al., 2024), our results highlight that enhanced multivariate decoding may be an equally important advantage. This suggests that OPM is not only valuable for precise localization but also for extracting distributed representational patterns that underpin modern cognitive neuroscience. Furthermore, the demonstrated robustness of OPM with reduced sensor density sets the stage for applying MVPA in populations such as infants and children, where small helmet sizes limit the number of sensors that can be accommodated. More broadly, the portability and flexibility of OPM systems open the door to studying cognition in more naturalistic and ecologically valid environments, and to extending brain–computer interface research to scenarios where lightweight and adaptable MEG solutions are needed. Taken together, these advances point toward a future where OPM-MEG expands both the methodological and conceptual horizons of neuroscience research.

## Acknowledgements

This work was supported by Wellcome Trust Discovery Award (grant number 227420), the NIHR Oxford Health Biomedical Research Centre (NIHR203316), Jiangsu Science and Technology Innovation Support Program (International Science and Technology Cooperation – Key National Industrial Technology R&D Cooperation Project SBZ2023000065) and Major Funding Project of the Jiangsu Industrial Technology Research Institute (JITRI).

The views expressed are those of the author(s) and not necessarily those of the NIHR or the Department of Health and Social Care. The Oxford University Centre for Integrative Neuroimaging was supported by core funding from the Wellcome Trust (203139/Z/16/Z and 203139/A/16/Z).

## CONFLICT OF INTEREST

The authors declare no conflicts of interest.

## DATA AVAILABILITY STATEMENT

Software used in this work has been properly cited within the manuscript. The code used for AMM with spherical harmonics is available on GitHub at https://github.com/Y-Bezs/AMM-with-spherical-harmonics.

## Notes

### Competing Interest Statement

The authors have declared no competing interest.

